# Molecular signatures of alternative fitness strategies in a facultatively social hover wasp

**DOI:** 10.1101/2022.12.02.518827

**Authors:** Benjamin A. Taylor, Daisy Taylor, Alexandrina Bodrug-Schepers, Francisco Câmara Ferreira, Nancy Stralis-Pavese, Heinz Himmelbauer, Roderic Guigó, Max Reuter, Seirian Sumner

## Abstract

Social insect queens and workers represent ideal models with which to understand the expression and regulation of alternative reproductive phenotypes. Most research in this area has focused on the molecular regulation of reproductive castes in obligately social taxa with complex social systems, while relatively few studies have addressed the molecular basis of caste in species in which the division of reproductive labour is more plastic. As a result, it is not clear whether, and to what extent, the mechanisms of caste in species with reproductive plasticity are the same as those that exist at the highest levels of social complexity. To address this knowledge gap, we analyse brain transcriptomic data for non-reproductives and reproductives of the facultatively social hover wasp Liostenogaster flavolineata, a representative of one of the simplest forms of social living. By experimentally manipulating the reproductive ‘queues’ exhibited by social groups of this species, we show that reproductive division of labour in this species is associated with surprisingly distinct transcriptomic signatures, similar to those observed in more complex social taxa; that variation in gene expression among non-reproductives reflects their investment into foraging effort more than their social rank; and that distinct co-expressed gene sets are associated with differential investment into alternative reproductive strategies. These results elucidate robust transcriptomic signals that represent the proximate basis of division of labour at the simplest level of insect sociality, and show these signals to be remarkably similar to those in more derived species.

## Introduction

Selection favours those organisms that are able to maximise the transmission of their genes to the next generation, but optimal strategies to achieve this transmission may vary considerably between ecological and social contexts, resulting in a range of alternative fitness strategies (Hazel et al. 2004). In many social species, for example, some individuals forego their own reproduction in favour of helping relatives. Such species are ideal models with which to determine the mechanisms that underlie alternative fitness strategies, since they exhibit a clear delineation between individuals that invest in either direct or indirect (‘altruistic’) fitness strategies. Understanding how and why indirect fitness strategies have evolved and are regulated has long been a major question in evolutionary biology.

In most social taxa, the evolution of indirect modes of reproduction can ultimately be explained in terms of inclusive fitness theory: altruistic behaviour evolves because altruists increase the fitness of individuals to whom they are related (Hamilton 1963; Foster et al. 2006). The question of how non-reproductive and reproductive phenotypes can arise from a shared genome at a proximate level is, however, less well-resolved. Research over the last two decades has begun to reveal how alternative reproductive strategies are produced through differential expression of shared molecular machinery (e.g. Colgan et al. 2011; Standage et al. 2016; Morandin et al. 2019), but most research in this area has focused on ‘obligately’ social insects that live in complex societies comprised of mutually dependent individuals, each of whom are committed to a specific role in the group (queen and worker ‘castes’). The molecular basis of the differentiation between castes in such obligately social species has been the focus of numerous studies, which have revealed that queens and workers exhibit strongly and consistently divergent patterns of gene expression (e.g. Morandin et al. 2015; He et al. 2019; Warner et al. 2019).

Despite the ecological importance of obligately social insect species (Wilson 1990), these species are relatively taxonomically limited. In insects, for example, such societies are found only in the ants, honey bees, stingless bees, bumble bees, vespine wasps and higher termites (Boomsma & Gawne 2017). Simpler forms of social organization, in which individuals retain the ability to switch between reproductive and non-reproductive roles within their lifetime, are much more common in both insects (e.g. polistine and stenograstrine wasps; halictid, xylocopid bees) and cooperatively breeding vertebrates (e.g. meerkats and long-tailed tits). Importantly, non-reproductive individuals in facultatively social species retain the ability to mate and to develop their reproductive physiology, and are potential successors to the dominant reproductive if she dies or if an alternative nesting opportunity arises (Field et al. 2006; Smith et al. 2009). Such individuals can be described as ‘hopeful reproductives’ that invest in altruistic behaviour whilst waiting for opportunities to reproduce (West-Eberhard 1975).

The flexible and reversible nature of their reproductive strategies makes facultatively social insects ideal models for understanding the emergence of the simplest forms of reproductive caste (Kronauer & Libbrecht 2018; Shell & Rehan 2018). For example, facultatively social bee taxa have been used to provide evidence for ‘molecular ground plan’ hypotheses (Kapheim et al. 2012, 2020), and to test the long-standing sociogenomic prediction that queen-biased genes should be relatively ancient and conserved compared to worker-biased genes (Jones et al. 2017). Genomic studies using facultatively social insects outside of the Anthophila (bees) are lacking, however. Accordingly, we have a limited and biased understanding of the proximate nature of alternative reproductive strategies in facultatively social insects, i.e. how the highly flexible alternative fitness strategies in these species are produced from a shared genome and how such plasticity is regulated (Simola et al. 2016; Libbrecht et al 2018; Taylor et al. 2021). Specifically, we would like to know whether the degree of molecular differentiation present among the uncommitted, transient castes of facultatively social species is similar to that which exists in obligately social species. Data regarding this question are largely absent outside of the Anthophila. In addition, few if any studies have addressed the question of the degree to which variation in individual-level investment in non-reproductive ‘worker’ activities is reflected at the molecular level. Species with reproductive ‘queues’ that predict investment into altruistic behaviours should be particularly suitable for the study of this topic.

Here we address these questions by conducting the first transcriptomic analyses for a facultatively social wasp, *Liostenogaster flavolineata*. This species represents the best-studied member of the stenogastrine hover wasps, a group which represents an independent evolutionary origin of facultative sociality (Bank et al. 2017), separate from all other social wasps. As in other facultatively social species, many *L. flavolineata* individuals spend part of their lives as non-reproductive ‘workers’, but such individuals may transition at some point to a reproductive ‘queen’ role, shifting between indirect and direct fitness strategies (Field et al. 2006; Bridge & Field 2007). *L. flavolineata* groups are small (2–10 individuals; Bridge & Field 2007) and consist of strictly age-determined dominance hierarchies, with the oldest non-reproductive individual in a group being first in line to replace the reproductive if she dies. Because they are more likely to attain a reproductive role in the near future, older (higher ranked) individuals exhibit a reduced foraging effort relative to younger (lower ranked) individuals, indicating a shift away from investment into risky behaviours as the chances of future reproductive opportunities improve (Field et al. 2006; Bridge & Field 2007). At any given time, however, only a single individual within a group acts as egg-layer, so *L. flavolineata* also exhibits the strong reproductive skew that is the defining characteristic of insect sociality (Sumner et al. 2002).

Based on these biological features, *L. flavolineata* represents an ideal model with which to address the outstanding questions outlined above and to shed light on the complex molecular origins of plastic social phenotypes. By sequencing the genome of this species, the first for any member of the Stenogastrinae, conducting controlled experiments to manipulate social hierarchy and caste behaviour, and examining in detail the influence of social rank and altruistic activity upon brain transcription, we generate novel insights into the molecular origins of plastic social phenotypes.

## Results

### L. flavolineata genome assembly

Illumina sequencing generated 163 million pairs of genomic PE reads and 102 million pairs of MP reads. The GC content distribution had a single peak both in the PE sequencing data that were used as input for the assembly and in the assembly itself, which contrasts with the bimodal or trimodal GC content distributions observed in other social hymenopterans (e.g. Kent et al. 2012; Harrop et al. 2020). The size of the genome estimated based on 17-mers was 373 Mbp. The assembled genome sequence obtained with SOAPdenovo was 291 Mbp, counting only contigs larger than 500bp. The longest scaffold in the assembly had a length of 5.23 Mbp, and the N50 scaffold length was 1.5 Mbp (**Supplementary Table S1**). Of 4,415 BUSCO groups searched, 97.9% were found in the assembly (96.9% complete). We therefore conclude that this assembly is a comprehensive representation of the *L. flavolineata* genome in terms of regions encoding gene sequences.

### *Phenotypic correlates of rank in* L. flavolineata

We first verified variation in caste-related phenotypes along the reproductive hierarchy by measuring ovarian development and time spent off-nest of individual wasps with different reproductive ranks. In unmanipulated colonies, there was a strongly significant negative relationship between within-group age rank and the amount of time individuals from unmanipulated groups spent off the nest (linear regression of time off-nest on rank: slope ± SE = -27.37 ± 1.95, *p* < 0.001, t_df_ = 78; **Figure 1A; Supplementary Table S2**).

**Figure 1.**
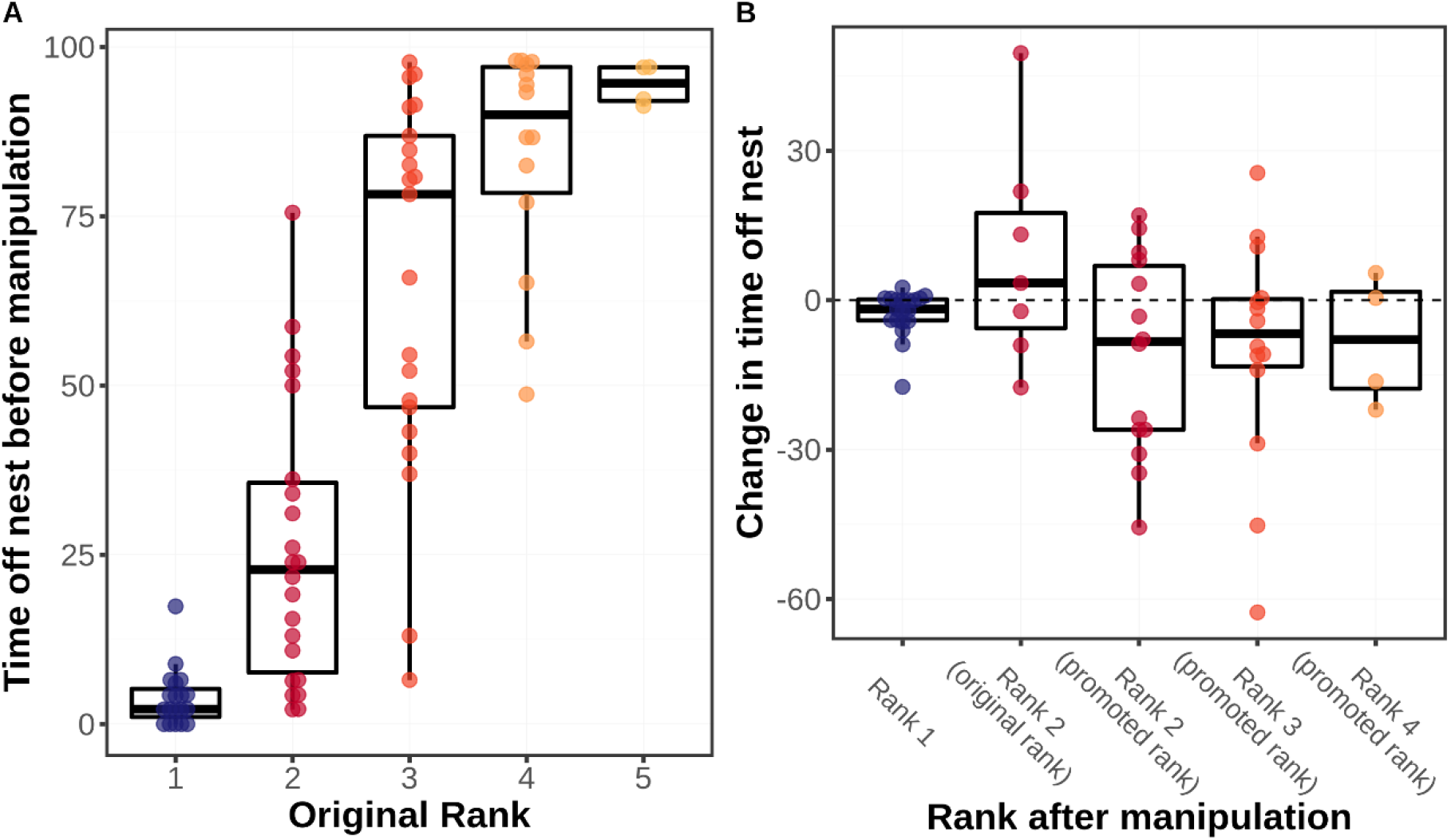
(A) Proportion of time spent off-nest before manipulation and (B) change in proportion of time spent off-nest following manipulation by individuals of different within-group ranks.

We also observed the expected changes in foraging efforts in response to manipulation of rank. Individuals that were promoted in rank by removing a higher-ranked individual reduced their time off the nest (and therefore putatively the time spent foraging) following manipulation (*n* = 35; two-sided Wilcoxon test W = 137, *p* = 0.017), although the dispersion of this change was high (mean ± SD = -10.16 ± 20.25; **Figure 1B**). By contrast, individuals whose rank was not manipulated showed no significant shift in their time spent off the nest (*n* = 26; mean ± SD = 0.35 ± 12.68; two-sided Wilcoxon W = 90, *p* = 0.2; **Figure 1B**), despite the fact that group size was decreased for these individuals following manipulation. This indicates that the removal of brood during manipulation was successful in maintaining a constant per-capita foraging requirement and supports the interpretation that behavioural changes in individuals with manipulated ranks were indeed due to their shift in the reproductive hierarchy.

Ovarian development was strongly dependent on rank at time of removal. The most dominant individual (Rank 1) within a nest was always inseminated and possessed several mature eggs in her ovarioles (*n* = 19; mean ± SD = 12.63 ± 2.29 eggs). By contrast, individuals of Rank 2 and below (*n* = 64) almost never possessed developed eggs at time of removal and were never found to be inseminated (**Supplementary Table S2**).

Overall, these results are in line with previous work (Field et al. 1999, 2006; Shreeves & Field 2002; Sumner et al. 2002; Bridge & Field 2007), showing that *L. flavolineata* groups are defined by a binary reproductive division of labour between individuals of Rank 1 (who are the sole egg-layers) and Rank 2 and beyond (non-reproductives). An age-based hierarchy exists among the non-reproductives, with reduced investment in foraging effort for higher-ranked individuals in the queue. We therefore posit that time spent off the nest can be taken as a proxy for time spent foraging and hereafter refer to the proportion of time spent off the nest by each individual as that individual’s ‘foraging effort’.

### Reproductive division of labour predicts gene expression variation

We next sequenced RNA from a number of focal individuals to identify patterns of gene expression associated with within-group rank. RNA was successfully sequenced from brain tissue taken from 83 individuals from 28 different nests, with an average of 14.4M successfully mapped reads per sample. Comparing all reproductives (Rank 1; *n* = 19) against all non-reproductives (Ranks 2–5; *n* = 64) with DESeq2, we identified 1117 differentially-expressed genes (DEGs). The 483 genes that were upregulated in reproductives (Rank 1) were associated with 19 significantly enriched GO terms (**Supplementary Tables S3-4**), including several terms related to DNA replication and histone binding. The 664 genes that were upregulated in non-reproductives (Ranks 2–5) were associated with 79 significantly enriched GO terms (**Supplementary Tables S5-6**), most of which related to respiration and metabolism.

We found strong signals of differential expression between Rank 1 individuals and individuals of each of the other ranks, with > 300 DEGs in each comparison (**Figure 2A**). This result held whether grouping individuals by their rank prior to manipulation or by their rank following manipulation (**Figure 2B**), indicating that manipulation did not seriously disrupt the reproductive division of labour within groups. The strongest differentiation in each case was between reproductive Rank 1 and Rank 3 individuals rather than the more worker-like Ranks 4 and 5. This is surprising, given that Rank 3 individuals exhibit intermediate foraging efforts (**Figure 1A**), but may be at least partially explained by the larger sample sizes for Rank 3 (pre-manipulation *n* = 21; post-manipulation *n* = 15) than Ranks 4 (pre-manipulation *n* = 15; post-manipulation *n* = 6) and 5 (pre-manipulation only; *n* = 6).

**Figure 2.**
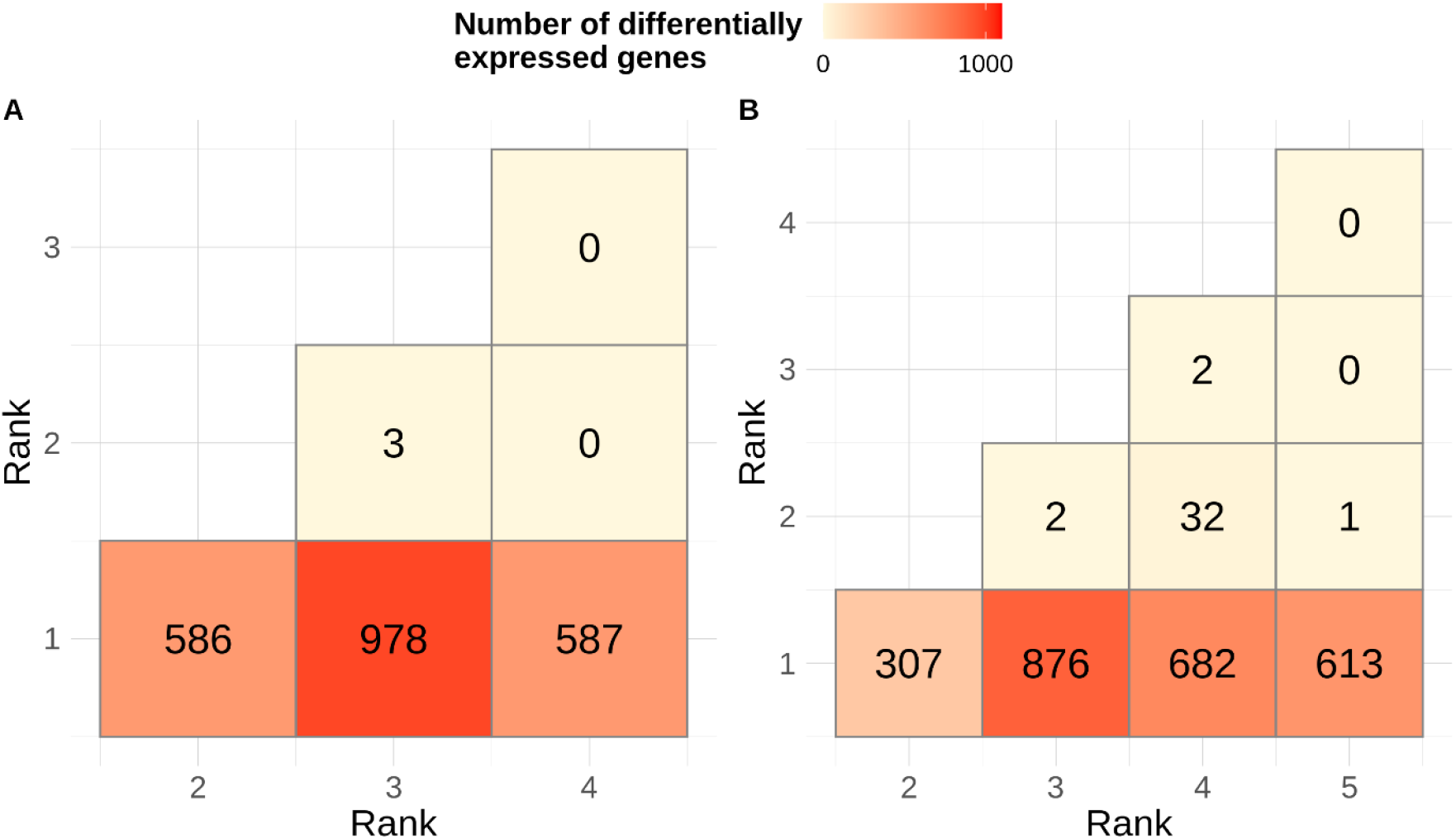
Number of genes differentially expressed between each pair of ranks. Ranks shown are those to which individuals belonged (A) after and (B) before manipulation. Post-manipulation rank sample sizes: R1 = 19; R2 = 21; R3 = 15; R4 = 6. Pre-manipulation rank sample sizes: R1 = 19; R2 = 22; R3 = 21; R4 = 15; R5 = 6.

### Effects of social hierarchy on brain transcription

#### Absolute foraging effort explains molecular differentiation among non-reproductives better than rank

We next focused on patterns of gene expression associated with the phenotypic variation observed within nests’ hierarchies. Over 1000 genes were identified as being differentially expressed with foraging effort and/or with rank, regardless of whether the rank considered was that identified before or after manipulation (**Table 2**). However, much of this differential expression was driven by reproductive individuals, which exhibit the highest rank and lowest foraging rates (**Figure 1A**). Excluding these individuals from the analysis and thereby focusing exclusively on non-reproductives, the number of DEGs identified with rank and/or foraging effort was 1–3 orders of magnitude lower (**Table 2**).

Analysis of these non-reproductive individuals (*n* = 64 wasps) revealed that foraging effort explains differences in gene expression more strongly than rank: 256 genes were correlated with foraging effort but only 45 genes with rank (**Table 2**). Furthermore, no genes were found to be differentially expressed with the residuals of rank on foraging effort, while 18 genes were differentially expressed with the residuals of foraging effort on rank, which suggests that it is foraging rate *per se* rather than rank that best predicts individual gene expression profiles. Surprisingly, there was no detectable effect of age itself on gene expression among non-reproductive individuals (**Table 2**), despite a substantial age range of 20 days between the oldest and youngest individuals in this analysis. Subsequent analyses therefore focused on the molecular signatures of foraging effort among non-reproductive individuals.

We next explored the functions of the genes (*n* = 256) that were putatively associated with foraging. Among non-reproductives, 173 genes were upregulated with respect to foraging rate. These were associated with 61 significantly enriched GO terms, many of which related to developmental and metabolic processes (**Supplementary Tables S7-8**). The 83 genes that were downregulated with respect to foraging rate were associated with 14 GO terms, including several terms related to metabolism or visual perception (**Supplementary Tables S9-10**).

Individuals that were promoted in rank after experimental removal of a higher-ranked wasp responded by modulating their foraging effort to match their new rank, and this change appears to have been matched at the level of gene expression. Individuals that were promoted from Rank 3 to Rank 2 exhibited a significant degree of gene expression differentiation when compared against Rank 3 individuals that had not changed their rank: 218 genes separated these two groups, and this set of DEGs overlapped significantly with those identified as being differentially expressed with foraging rate among non-reproductives (two-sided hypergeometric test *p* = 0.016; **Supplementary Table S11**). Meanwhile, individuals that were promoted to Rank 2 showed little differentiation from Rank 2 individuals that had not changed their rank (10 DEGs; **Supplementary Table S12**). These results suggest that the shifts in investment in individual-level foraging effort that accompany a change in rank are also reflected at the level of the brain transcriptome, with individuals adopting a transcriptional profile that is closer to their new rank than to the rank that they possessed prior to promotion.

#### Distinct suites of co-expressed genes are associated with foraging effort

Across the brain transcriptomes of all non-reproductive individuals (*n* = 64), we identified 20 distinct co-regulated gene modules ranging in size from 32 to 1831 genes. None of these modules showed expression levels correlated with rank (either before or after manipulation) nor with age, but two exhibited strong (but opposing) correlations with foraging effort (**Figure 3**).

**Figure 3.**
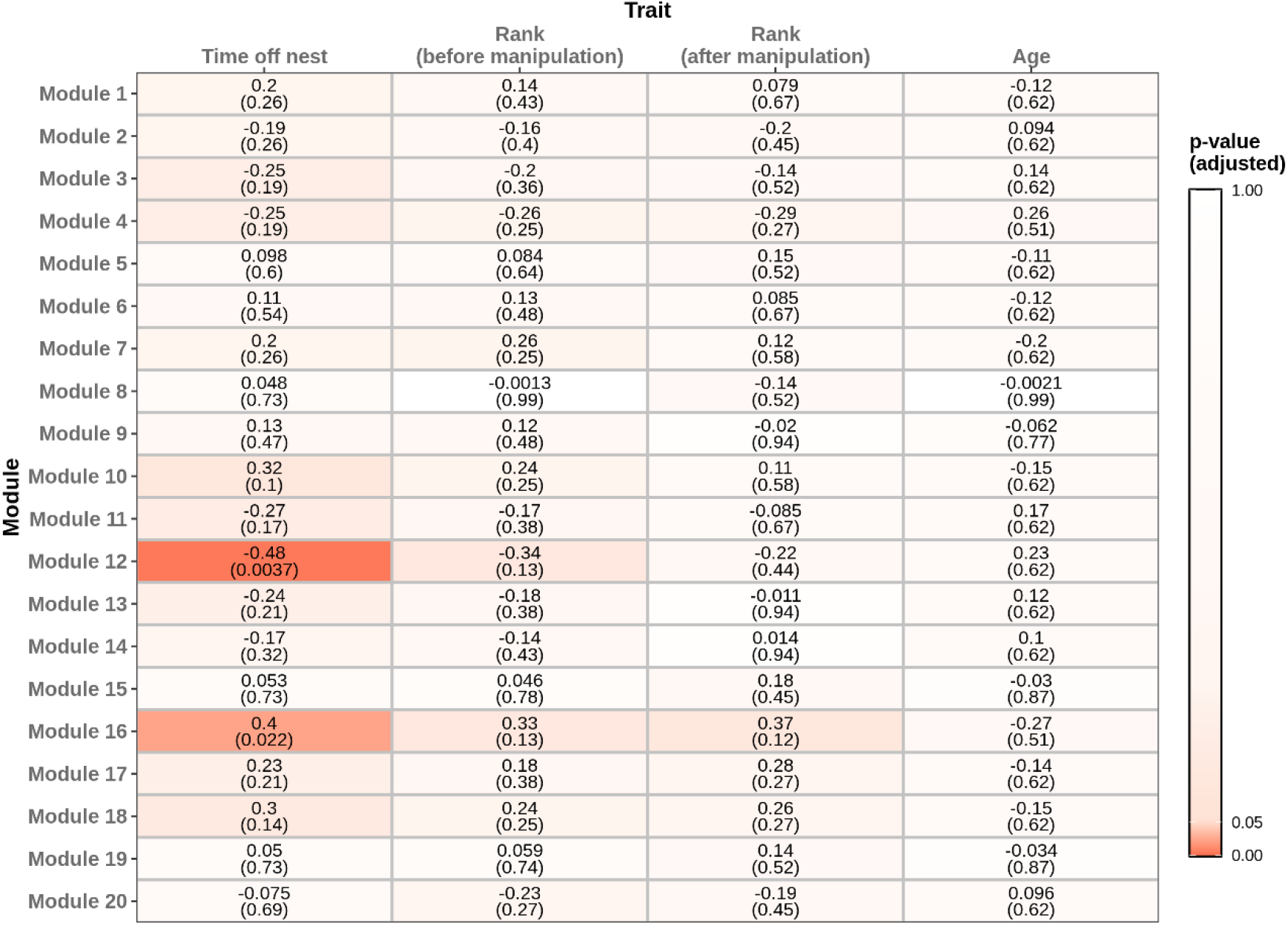
Association of phenotypic traits (time off nest, which is a proxy for foraging effort; rank assessed before manipulation; rank assessed after manipulation; and age) among non-reproductive individuals with co-expressed gene modules present in those individuals. Number outside parentheses: Pearson correlation. Number within parentheses: FDR-corrected *p*-value.

One module (Module 12; *n* = 83 genes) was significantly negatively associated with foraging rate (*r* = -0.48, *p* = 0.004). Genes that were more strongly correlated with foraging rate, whether positively or negatively, were also more strongly correlated with the eigengene of Module 12 (i.e. they had higher ‘module membership’), suggesting a meaningful relationship between the expression of this module and individuals’ foraging effort (**Figure 4A**). Furthermore, genes that were part of this module overlapped significantly both with genes that were found to be negatively correlated with respect to foraging rate when using DESeq2 (two-sided hypergeometric test *p* < 0.001), and also with genes that were reproductive-biased (i.e. those that were upregulated in reproductives versus non-reproductives; two-sided hypergeometric test *p* = 0.016). However, we did not find this same pattern of overlap at the level of putative gene function. The 83 genes contained within Module 12 were enriched for 17 GO terms, several of them associated with visual processing (**Supplementary Tables S13-14**), but these terms did not overlap significantly with those that were reproductive-biased (two-sided hypergeometric test *p* = 1), nor with those that were negatively correlated with foraging rate (two-sided hypergeometric test *p* = 0.116).

**Figure 4.**
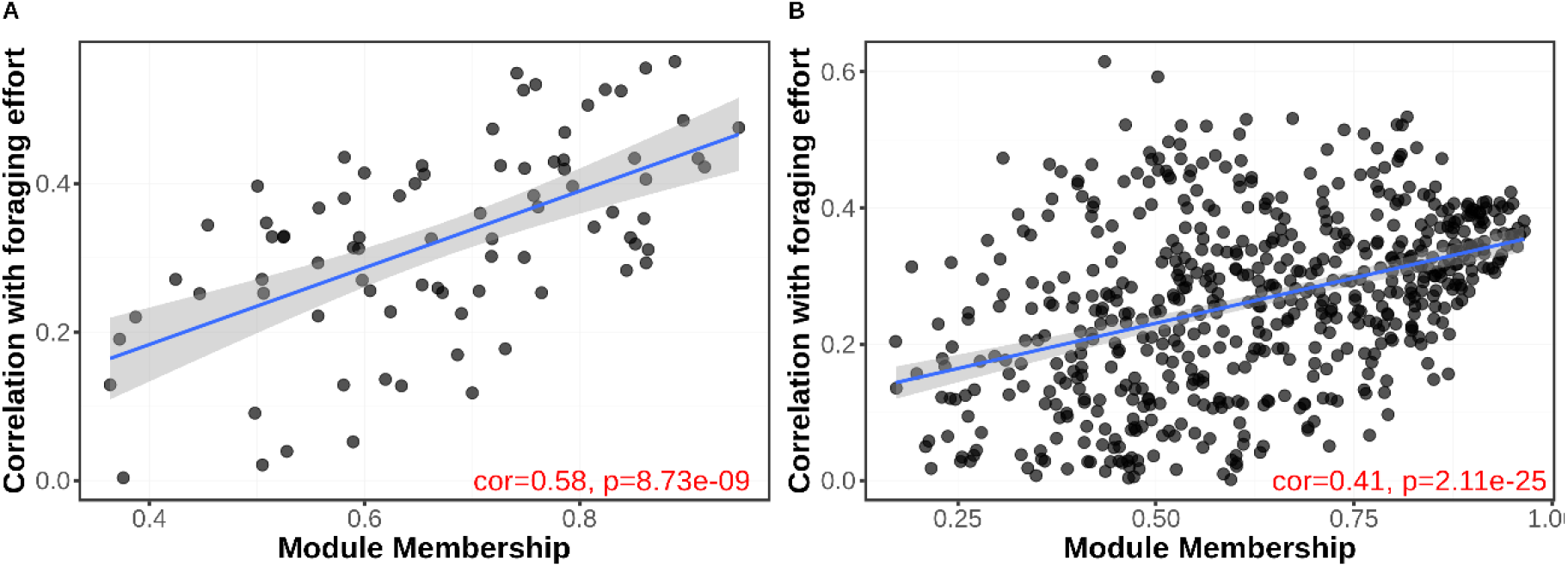
Strength of absolute (unsigned) correlation of genes with foraging effort among non-reproductives, plotted against module membership of those genes within (A) Module 12 and (B) Module 16.

A second, larger module (Module 16; *n* = 606 genes) exhibited a strongly positive association with foraging rate (*r* = 0.4, *p* = 0.022). Genes that were more strongly associated with foraging effort were also more strongly correlated with this module’s eigengene (**Figure 4B**). The genes in this module were associated with 94 GO terms, including many terms associated with respiration, and metabolic and biosynthetic processes (**Supplementary Tables S15-16**). Contrary to Module 12, both genes and GO terms associated with Module 16 overlapped significantly with the set of genes and terms that were upregulated with foraging rate among non-reproductives, as measured by DESeq2 (genes: two-sided hypergeometric test *p* < 0.001; GO terms: two-sided hypergeometric test *p* < 0.001), and genes and GO terms that were upregulated in non-reproductives versus reproductives (genes: two-sided hypergeometric test *p* < 0.001; GO terms: two-sided hypergeometric test *p* < 0.001).

## Discussion

Facultatively social insects are a valuable model with which to study the evolution of plastically-expressed alternative fitness strategies. Until recently, however, their potential to shed light on the mechanisms associated with such strategies has not been fully exploited, as most studies have concentrated on caste differences in obligately social species. Here, we have addressed this knowledge gap by examining the brain gene expression profiles associated with social organization in reproductively skewed groups of the facultatively social hover wasp *L. flavolineata*.

Caste differences in obligately social insect species are stable and associated with strong signatures of molecular differentiation (e.g. Morandin et al. 2015; He et al. 2019; Warner et al. 2019). Whether such strong patterns of molecular differentiation are also present between reproductives and non-reproductives in species that exhibit a high degree of social plasticity is less clear, especially outside the few facultatively social bee species that have been studied. Our results indicate that patterns of molecular differentiation in the social groups of *L. flavolineata*, a facultatively social wasp that represents an independent and little-studied origin of sociality, substantially resemble those found in the groups of obligately social insects. In this species, as in more socially complex taxa, the strongest signatures of within-group differentiation are found between reproductives and non-reproductives, despite the presence of significant behavioural variation among non-reproductives. That we were able to detect such large differences in brain transcriptomes may seem surprising given that the behaviour of Rank 2 individuals in this species is intermediate between that of reproductives (Rank 1 individuals) which do not forage at all (Field & Foster 1999; Cant & Field 2001; Shreeves & Field 2002), and low-ranked non-reproductives which perform the majority of the group’s foraging (**Figure 1A**). However, this finding is in agreement with a number of studies that have indicated a strong relationship between ovarian activation and brain gene expression (Amdam et al. 2006; Wang et al. 2009, 2010), and reinforces the notion that ovarian activation is the defining characteristic of division of labour in social insects.

Focusing on transcriptomic differences among non-reproductives, we identified approximately 250 genes whose expression varied in line with foraging rate—a substantial number, though significantly smaller than that associated with caste differences. A plausible null hypothesis is that this expression variation simply reflects phenotypic variation among individuals; for example, differential expression with foraging rate might stem from the energetic demands of spending more time in flight. However, we believe that three factors speak against this null hypothesis and in favour of the alternative hypothesis that gene expression variation among non-reproductives reflects variation in investment into alternative fitness strategies.

First, if within-group gene expression differences primarily reflect differences in e.g. energetic expenditure, then we should have observed greater differences between Rank 2 individuals and Rank 4/Rank 5 individuals than between Rank 2 individuals and Rank 1 reproductives, who are more similar in foraging effort (**Figure 1A**), yet this was not the case (**Figure 2**). Second, individuals that were promoted from Rank 3 to Rank 2 shifted gene expression to match their new rank, which strongly suggests that gene expression variation is not primarily structured by independent factors such as age. Third, the majority of the GO terms that were enriched among foraging-biased genes (both positively and negatively) related to relatively basal metabolic and developmental processes, a pattern similar to that observed in caste-biased GO terms in both *L. flavolineata* and the paper wasp *Polistes dominula* (Standage et al. 2016; Taylor et al. 2021). This seems to belie the possibility that these genes were solely associated with such specific behaviours as e.g. flying and foraging. In fact, GO terms for visual processing were enriched among those genes that were negatively associated with foraging rate, a surprising result given that foragers presumably rely on visual signals to locate prey. Although we acknowledge that GO annotations are frequently patchy and incomplete, overall these factors support the hypothesis that gene expression variation among *L. flavolineata* non-reproductives genuinely represents variation in individuals’ potential fitness (i.e. their likelihood of achieving a reproductive position in the near future), over and above simply reflecting differences in age or energy expenditure.

Although we hypothesise that it is prospective reproductive opportunities rather than energetic expenditure alone that explains the variation in gene expression observed among non-reproductives, we identified a larger number of genes differentially expressed with foraging rate than with rank. Moreover, the set of rank-biased genes was almost entirely subsumed by the set of foraging-biased genes. This may indicate that rank is in fact a relatively crude indicator of individual-level variation in proximity to the reproductive role. Within-rank variation in the foraging rate of Rank 2 and Rank 3 individuals was very high relative to that of e.g. Rank 1 individuals (**Figure 1A**), which suggests that factors other than rank influence individuals’ foraging effort. Nest size is known to be one such factor (Shreeves & Field 2002; Cant & English 2006), but others might include the age, health and fecundity of the incumbent reproductive (Kokko & Johnstone 1999; Bridge & Field 2007). If individuals modulate their foraging rate based on the likelihood of future inheritance of the reproductive position, and that likelihood is affected by e.g. the projected mortality of the current reproductive, then observed foraging rate might actually be a stronger proxy of individual-level alternative fitness strategies than rank itself.

Facultatively social insects are important models with which to investigate the earliest stages of insect social evolution, in which individuals can switch opportunistically between direct and indirect fitness strategies. Social groups in the facultatively social hover wasp *L. flavolineata* are characterised by linear hierarchies, and an individual’s position in the resulting ‘queue’ dictates her investment into altruistic foraging behaviour, a system that lends itself well to studies of the molecular correlates of alternative fitness strategies. In this study, we have shown that gene expression in *L. flavolineata* colonies is structured by both actualised and potential direct fitness, that individuals are able to facultatively shift their gene expression profiles to match changes in direct fitness prospects, and that distinct co-expression modules are associated with alternative fitness strategies. These results represent the first in-depth study of the molecular basis of social behaviour in a vespid species outside of the Polistinae, the only such analysis for a facultatively social wasp, and the first genome sequence for a Stenogastrine wasp, representing an independent evolutionary lineage of insect sociality. These resources and the insights provided by our analyses open fruitful lines for future research into the maintenance and evolution of sociality, and reveal robustly co-expressed signals of transcriptomic differentiation that appear to track differential investment into reproductive versus non-reproductive behavioural strategies.

## Materials and Methods

### Field monitoring and behavioural experiments

#### Experimental set-up

Fieldwork was undertaken in Fraser’s Hill, Malaysia between January and April 2017. *L. flavolineata* nests from aggregations situated in under-road culverts were selected for study on the basis of the presence of multiple pupal caps and observation of active egg-laying. Over 2 days, all individuals on each nest were given a unique combination of coloured paint marks to facilitate subsequent individual-level identification. Confirmation that all wasps had been successfully marked was achieved by censuses of group members at night, when all individuals are present on the nest. Brood were mapped in each nest to confirm the presence of an egg-layer, and to identify pupae which would shortly hatch. Reproductives were identified by observation of egg-laying or through censuses to determine foraging effort: reproductives rarely leave the nest in this species (Field & Foster 1999; Cant & Field 2001; Shreeves & Field 2002). Nests were monitored every day to measure foraging effort and identify newly emerging wasps. Newly emerged group members were identified by the co-appearance of an unmarked wasp and a hatched pupal cell. Once identified, newly emerged individuals were left on the nest for 3 days before marking, to avoid interfering with their nest orientation flights.

Focal nests (*n* = 28) consisting of 3–5 wasps of known age and rank were generated by removing unmarked wasps and/or wasps of unknown age; such group sizes are typical of natural nests in this species (Field et al. 2000; Shreeves & Field 2002). To achieve this, nests were censused daily until 2–4 new individuals had emerged. These newly emerged individuals, together with the nest’s reproductive (Rank 1), were used as the focal individuals for the rest of the study. To generate nests of comparable group sizes and with non-reproductives of known ages, all other wasps were removed at dawn on the day following the emergence of the requisite number of focal wasps. For 10 days following this manipulation, nests were censused every ∼30 minutes during peak foraging hours (07:00–11:00) to quantify time spent off the nest for each individual. Time spent off the nest is thought to be a reliable proxy for foraging effort in *L. flavolineata* (Field & Cronin 2006; Bridge & Field 2007): older individuals are more likely to inherit the position of egg layer and are therefore expected to invest less in risky foraging behaviour (Field & Cronin 2006; Bridge & Field 2007). From this initial manipulation until the end of the experiment, additional wasps that emerged from the nest were removed to ensure that colony sizes remained constant. Thus, at the end of the 10-day period, the rank and average time spent off-nest were known for each individual from 28 focal nests consisting of a single reproductive (exact age unknown but >2 months old) and 2–4 non-reproductives of known ages (within-rank age range ≈ 5 days). These preliminary manipulations allowed us to generate a starting population of nests for which factors such as age, group size, brood number and foraging effort were quantified and standardised for use in our experiment and analyses.

#### Experiments and sampling

Using an experimental design based on that employed by Field et al. (2006), we performed manipulations in which the second-ranked (Rank 2) or third-ranked (Rank 3) individual from each nest was removed in order to promote lower-ranked wasps to a higher rank within the group hierarchy. To promote non-reproductive wasps on each nest either from Rank 3 to Rank 2 or from Rank 4 to Rank 3, at dawn on day 11, a single focal wasp was removed from each of the 28 nests and placed directly into RNAlater (Thermo Fisher Scientific). The removed individual was always of either Rank 2 (*n* = 15; age mean ± SD = 30.3 ± 3.1 days) or Rank 3 (*n* = 7; age mean ± SD = 25.3 ± 1.4 days). To ensure that the ratio of helpers to brood remained constant despite the loss of a helper, brood were also removed from the nests at this time. Brood was divided into three categories: eggs, small/medium larvae and large larvae. A proportion, R/N (where R is the number of adults removed and N is the original number of wasps), of each category was removed using fine tweezers. Pupae, which do not require feeding, were not removed. Nests were given 48 hours to settle, after which censuses were performed daily for 5 days using the same methodology as previously. At the end of the 5-day censusing period, all wasps were removed from the nest before dawn: individuals’ heads were placed directly into RNAlater for gene expression analysis, and their bodies into 95% EtOH for dissection. The ages of non-reproductives of the same final rank collected at this stage that were subsequently sequenced were approximately age-matched across nests (age mean ± SD: Rank 2 = 33.3 ± 4.3 days; Rank 3 = 27.0 ± 1.6 days; Rank 4 = 22.8 ± 1.5 days). The ovarioles of each individual were dissected and the number of developed eggs counted. The mating status of each individual was assessed by examining the spermatheca for the presence of sperm.

This design allowed us to compare gene expression between reproductives (Rank 1) and non-reproductives (Ranks 2–5). The design also enabled us to assess the extent to which variation in gene expression varied among non-reproductives of different ranks (Ranks 2–5) and, therefore, with investment into risky foraging behaviour. If rank and/or foraging effort are reflected at the level of brain transcription, then we expected that fine-scale molecular differentiation among non-reproductive ranks (i.e. Ranks 2–5) might be as profound as that between reproductive castes (i.e. Rank 1 versus Ranks 2–5). Finally, our chosen experimental design allowed us to compare the brain gene expression patterns of individuals that had been promoted in rank to those which had not undergone promotion. We predicted that individuals that had been promoted from Rank 3 to Rank 2 would exhibit a concomitant shift in rank- and foraging-related gene expression: if this were the case, then individuals promoted from Rank 3 to Rank 2 should more closely match the expression patterns of unmanipulated Rank 2 individuals than of unmanipulated Rank 3 individuals.

### Genomic and transcriptomic analyses

#### Genome sequencing and assembly

Complete methods and results for the assembly are given in **Supplementary Document S1**. In brief: DNA was extracted from a single haploid *L. flavolineata* male using the DNeasy Blood & Tissue Kit (Qiagen) according to the manufacturer’s instructions. DNA quantification was performed with a Qubit 3.0 fluorometer using the dsDNA BR assay kit (Thermo Fisher, Waltham, MA, USA) and DNA integrity was monitored on an agarose gel. The preparation of sequencing libraries was performed using DNA isolated from haploid males. A paired-end (PE) sequencing library with a peak insert size of 535 bp was constructed using 200 ng of genomic DNA with a TruSeq Nano LT library preparation kit (Cat # FC-121-4002, Illumina, San Diego, CA, USA) according to the kit supplier’s instructions. A mate-pair (MP) library with a peak span size of 1450 bp and a mean span size of 1027 bp was prepared from 570 ng of genomic DNA by tagmentation using an MP library preparation kit (Cat # FC-132-1001, Illumina, San Diego, CA, USA), without size selection. The MP library was amplified using 12 cycles of PCR. The quality and quantity of the libraries was checked on a DNA 1000 chip on the Agilent Bioanalyzer 2100 (Agilent, Santa Clara, CA, USA).

Illumina sequencing was performed on a HiSeq 2500 instrument utilising v4 Illumina sequencing chemistry, combined with a 2×125 cycle sequencing recipe. Raw sequencing data underwent quality control with FastQC (Andrews 2010), thereafter, trimmomatic (Bolger et al. 2014) was employed for data filtering based on phred scores, using the following parameters: LEADING:25 TRAILING:25 SLIDINGWINDOW:10:25 MINLEN:36. Genome assemblies were preformed using SOAPdenovo_v2.04 (Luo et al. 2012). Pre-assemblies were first calculated based on PE reads which were assembled either as single reads or as pairs (Dohm et al. 2014), in order to assess the insert size distribution (PE reads) and span size distribution (MP reads) of the sequencing libraries, respectively. Bowtie2 (Langmead & Salzberg 2012) was used with an insert size interval between 100 and 1200 and 1 million PE read-pairs were sampled to estimate the library insert size. To estimate the span size of the MP read-pairs, bowtie2 was used on an assembly version using the paired-end library as pairs and an insert size interval between 100 and 20000, and sampled 1 million MP read-pairs. Using the determined library insert size and MP span size as parameters for the assembly run, several assemblies were calculated from the quality-filtered sequencing reads by varying the k-mer size parameter between 23 and 125. An assembly calculated with k-mer size 69 was the best performing in terms of assembly metrics as assessed by QUAST (Gurevitch et al. 2013).

BUSCOv3 (Simão et al. 2015; Waterhouse et al. 2018) was used in genome mode with the hymenoptera_odb9 lineage and honeybee1 species to assess assembly completeness (blast 2.2.30, AUGUSTUS 3.2.1). Metrics of the final assembly were determined with custom scripts, taking only sequences larger than 500 bp into account. Jellyfish 2.2.10 (Marçais & Kingsford 2011) was used to determine genome size based on the quality-filtered Illumina PE sequencing reads. Bioawk was used to retrieve the GC content of all the reads as well as non-overlapping 125 nt segments of the final assembly lacking undetermined bases (no unknown nucleotide “N”).

#### Gene expression quantification

Brain tissue was extracted from the heads of a subset of individual focal samples and RNA was extracted using the RNeasy Mini Kit (Qiagen) according to the manufacturer’s instructions. Library preparation was performed by Novogene Co. followed by sequencing on an Illumina HiSeq 2000 platform with 150-bp paired-end reads to a depth of 30 million reads/sample. Transcript filtering, trimming, alignment and quantification were performed with the aid of Nextflow using the default options provided by the nf-core/rnaseq pipeline v1.4.2 (Ewels et al. 2020): adapter and quality trimming were performed using TrimGalore (Kreuger 2015), followed by removal of ribosomal sequences using SortMeRNA (Kopylova et al. 2012); read alignment against the *L. flavolineata* genome was performed using STAR (Dobin et al. 2013) followed by quantification using Salmon (Patro et al. 2017); aligned reads were assembled into genes using StringTie2 (Kovaka et al. 2019). Quality control (QC) checks identified six samples (three Rank 1s; two Rank 2s; one Rank 4) as exhibiting low sequencing quality, and these samples were excluded from subsequent analyses as a result. Finally, read counts were subjected to a round of filtering to remove any gene that was not expressed at a minimum of one count/sample in at least one set of wasps (as grouped by original rank). Following this filtering, 11258/14095 (79.9%) genes remained.

#### Differential expression analysis

Differential expression analyses were performed in R using the DESeq2 package (Love et al. 2014). Unless otherwise stated, differential expression was calculated relative to a baseline fold change of 0. Where a fold change threshold was used, this value became the baseline against which differential expression was measured (i.e. for a fold change threshold of 1.5, genes were considered differentially expressed if their absolute fold change was significantly greater than 1.5). Genes were considered differentially expressed between conditions if *p* < 0.05 after false discovery rate correction according to the Benjamini-Hochberg procedure.

#### Gene ontology (GO) enrichment analysis

To perform GO enrichment analysis, we first identified reciprocal BLAST best hits between *L. flavolineata* and *Drosophila melanogaster* proteins using BLAST+ (Camacho et al. 2006). 8266/11258 (73.4%) *L. flavolineata* genes possessed a reciprocal best hit with *D. melanogaster*. GO annotations for each D. melanogaster gene were acquired from BioMart (Smedley et al. 2009) and each *L. flavolineata* gene was assigned the GO terms of its reciprocal best hit. GO enrichment analysis was then performed in R via the topGO package (Alexa & Rahnenfuhrer 2009) using TopGO’s weight01 algorithm and Fisher’s exact test to identify GO terms that were significantly overrepresented (*p* < 0.01) in a focal set of genes against a background consisting of all genes that appeared in the relevant analysis.

#### Gene co-expression network analysis

We sought to identify co-expressed modules of genes associated with correlates of risky foraging activity among non-reproductives. To achieve this, weighted gene co-expression network analysis was performed in R using the WGCNA package (Langfelder & Horvath 2008). As WGCNA is particularly sensitive to genes with low expression, data were first subjected to a second round of filtering in which genes that had < 10 reads in > 90% of sampled individuals were removed, as recommended by the package authors. This second round of filtering removed an additional 1821 genes, leaving a total of 9428 genes. Counts were subjected to DESeq2’s variance-stabilizing transformation prior to further analysis. Consensus gene modules across all non-reproductives were then constructed using a soft-threshold power of 6 and the signed hybrid adjacency criterion. Network summary measures and gene dendrograms for this analysis are provided in **Supplementary Figures S1-2**. Initially, 26 gene modules were identified. Modules whose eigengene correlation was > 75% were subsequently merged, after which 20 consensus modules remained. Finally, the Pearson correlation of each module eigengene with each phenotypic trait was calculated and subjected to Benjamini-Hochberg FDR correction.

## Acknowledgments

We would like to thank S. Durai and R. Hashim for logistical support, and E. Favreau and O. Corbett for analytic advice and comments on an early version of the manuscript. Genomic sequencing was performed at the Vienna BioCenter Core Facilities (VBCF), Vienna, Austria. This work was conducted under research and collecting permits from the Malaysian Economic Planning Unit (UPE 40/200/19/3379). This work was supported by the Natural Environment Research Council (NE/L002485/1 awarded to BAT and NE/ M012913/2 awarded to SS) and by the Biotechnology and Biological Sciences Research Council (BB/R003882/1 and BB/S003681/1 awarded to MR).

## Figures and Tables

**Table 1.**
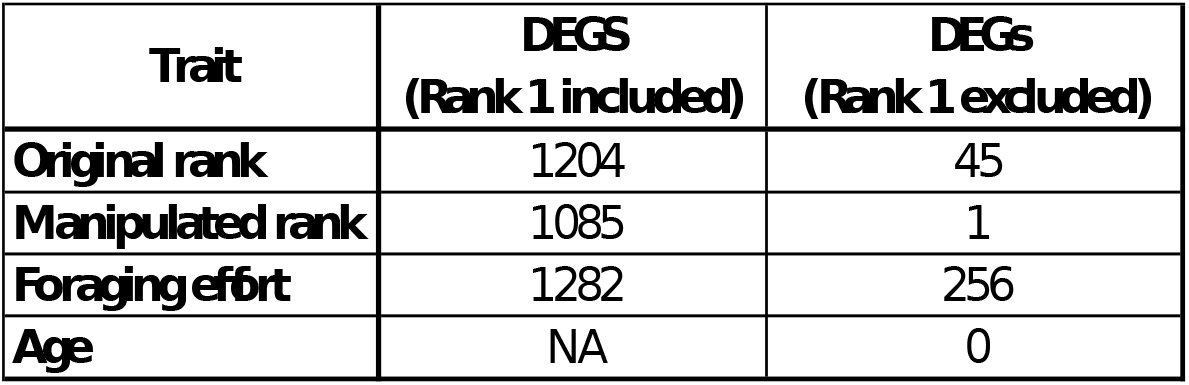
Number of differentially-expressed genes (DEGs) whose expression was correlated with continuous traits across all individuals or when excluding Rank 1 individuals. Age data were not available for Rank 1 individuals and so it was not possible to identify genes differentially expressed with age when including this group; however, no effect of age was detected for the set of wasps of known ages. Pre-manipulation *n* = 64 non-reproductive and 19 reproductive wasps; post-manipulation *n* = 42 non-reproductive and 19 reproductive wasps.

| **Trait** | **DEGS**<br>**(Rank 1 included)** | **DEGS**<br>**(Rank 1 excluded)** |
| --- | --- | --- |
| **Original rank** | 1204 | 45 |
| **Manipulated rank** | 1085 | 1 |
| **Foraging effort** | 1282 | 256 |
| **Age** | NA | 0 |

