## Supplemental Document S1 for "Molecular signatures of alternative fitness strategies in a facultatively social hover wasp"

### Supplementary Document S1 – *Liostenogaster flavolineata* genome assembly

#### Methods

DNA was extracted from a single haploid *L. flavolineata* male using a DNeasy Blood & Tissue Kit (Qiagen) according to the manufacturers' instructions. DNA quantification was performed with a Qubit 3.0 fluorometer using a dsDNA BR assay kit (Thermo Fisher, Waltham, MA, USA) and DNA integrity was monitored on an agarose gel. The preparation of sequencing libraries was performed using DNA isolated from haploid males. A sequencing library with a peak insert size of 535 bp was constructed using 200 ng of genomic DNA with a TruSeq Nano LT library preparation kit (Cat # FC-121-4002, Illumina, San Diego, CA, USA) according to the kit supplier´s instructions. A mate-pair (MP) library with a peak span size of 1450 bp and a mean span size of 1027 bp was prepared from 570 ng of genomic DNA by tagmentation using an MP library preparation kit (Cat # FC-132-1001, Illumina, San Diego, CA, USA), without size selection. The MP library was amplified using 12 cycles of PCR. The quality and quantity of the libraries was checked on a DNA 1000 chip on the Agilent Bioanalyzer 2100 (Agilent, Santa Clara, CA, USA).

Illumina sequencing was performed at the Vienna BioCenter Core Facilities (VBCF), Vienna, Austria, on a HiSeq 2500 instrument utilising v4 Illumina sequencing chemistry, combined with a 2x125 cycle sequencing recipe. Raw sequencing data underwent quality control with FastQC (Andrews 2010), thereafter, trimmomatic (Bolger et al. 2014) was employed for data filtering based on phred scores, using the following parameters: LEADING:25 TRAILING:25 SLIDINGWINDOW:10:25 MINLEN:36. Genome assemblies were preformed using SOAPdenovo_v2.04 (Luo et al. 2012). Pre-assemblies were first calculated based on paired-end (PE) reads which were assembled either as single reads or as PE reads (Dohm et al. 2014), in order to assess the insert size distribution (PE reads) and span size distribution (MP reads) of the sequencing libraries, respectively. Bowtie2 (Langmead & Salzberg 2012) was used with an insert size interval between 100 and 1200 (bowtie2 --fr -I 100 -X 1200 -1 pe_reads1.fastq -2 pe_reads2.fastq -x preassembly_1) and 1 million PE read-pairs were sampled to estimate the library insert size. To estimate the span size of the MP read-pairs, bowtie2 was used on an assembly version using the paired-end library as pairs and an insert size interval between 100 and 20000 (bowtie2 --rf -I 100 -X 20000 -1 mp_reads1.fastq -2 mp_reads2.fastq -x preassembly_2), and sampled 1 million MPs. Using the determined library insert size and MP span size as parameters for the assembly run, several assemblies were calculated from the quality-filtered sequencing reads by varying the k-mer size parameter between 23 and 125. An assembly calculated with k-mer size 69 (SOAPdenovo-63mer all -s assembly.config -K 69 -R -o assembly.K69) was the best performing in terms of assembly metrics as assessed by QUAST (Gurevitch et al. 2013).

BUSCOv3 (Simão et al. 2015; Waterhouse et al. 2017) was used in genome mode with the hymenoptera_odb9 lineage and honeybee1 species to assess assembly completeness (blast 2.2.30, AUGUSTUS 3.2.1). Metrics of the final assembly were determined with custom scripts, taking only sequences larger than 500 bp into account. Jellyfish 2.2.10 (Marçais & Kingsford 2011) was used to determine genome size based on the quality-filtered Illumina PE sequencing reads. Bioawk was used to retrieve the GC content of all the reads as well as non-overlapping 125 nt segments of the final assembly lacking undetermined bases (no unknown nucleotide "N").

The genome of *L. flavolineata* was assembled from DNA extracted from a single haploid male. From this individual, both a paired-end sequencing library and a mate-pair library were prepared (see methods for details). By means of Illumina sequencing,163 million pairs of genomic paired-end reads and 102 million pairs of mate-pairs were obtained. The size of the genome was estimated by counting k-mers, using the unassembled paired-end sequencing data as input for an analysis using jellyfish (Marçais & Kingsford 2011). In this way, a genome size of 373 Mbp was calculated for *L. flavolineata* based on 17-mers. The genome sequence that was assembled using SOAPdenovo_v2.04 was smaller, i.e. 291 Mbp, taking account only of sequences > 500 bp. The longest scaffold in the *L. flavolineata* assembly Lifl-v1.0 had a length of 5.22 Mbp, and the N50 scaffold length was 1.5 Mbp (**Table SA1**). A fraction of the total assembly was contained within sequences <= 500 nt, i.e. 243,289 sequences (30 Mbp). The GC content distribution had a single peak in both in the PE sequencing data that were used as input for the assembly and the assembly itself, which contrasts with findings of bimodal or trimodal GC content distributions in other wasps. The completeness of the *L. flavolineata* genome assembly was assessed with respect to conserved hymenopteran genes, using the Benchmarking Universal Single-Copy Orthologs (BUSCO) approach (Simão et al. 2015; Waterhouse et al. 2017). Of 4,415 BUSCO groups searched, 97.9% were found in the assembly, and 96.9% were complete (**Table SA2**). We may therefore conclude that the Lifl-v1.0 genome assembly is a highly comprehensive representation of the *L. flavolineata* genome.

| | |
|---|---|
| **Assembly size** | 291.28 Mbp |
| **N50 size** | 1.50 Mbp |
| **% GC** | 41.65 |
| **% unspecified bases (N)** | 5.1 |
| **Largest scaffold** | 5.22 Mbp |
| **Number of scaffolds + contigs** | 3,541 |

**Table SA1.** *L. flavolineata* genome assembly metrics based on sequences > 500 bp.

| BUSCO category | Number | Percentage |
|---|---|---|
| **Complete BUSCOs** | 4,277 | 96.9 |
| **Complete Single-Copy BUSCOs** | 4,268 | 96.7 |
| **Complete Duplicated BUSCOs** | 9 | 0.2 |
| **Fragmented BUSCOs** | 46 | 1 |
| **Missing BUSCOs** | 92 | 2.1 |
| **Total BUSCO number of groups** | 4,415 | 100 |

**Table SA2.** BUSCO completeness metrics for the *L. flavolineata* genome assembly.
