## Supplemental Document S2 for "Molecular signatures of alternative fitness strategies in a facultatively social hover wasp"

### **Supplementary Document S2 – *Liostenogaster flavolineata* genome structural annotation**

#### Repeat masking of the *L. flavolineata* genome assembly

The genome assembly was first modified by removing all contigs with fewer than 500 bases as these were previously determined to most likely not to include any complete genes using the genome assembly "gene space" assessment program BUSCOv2 (Simão et al. 2015). Then we executed RepeatMasker (v4.0.5) using hymenopteran repeat libraries in order to find and mask transposable elements as these reduce the accuracy of gene prediction tools. We used RepeatMasker through the repeat masking module of the genome annotation program Maker v2.31.9 (Campbell et al. 2014) which also uses RepeatRunner to further increase the accuracy of transposon finding. We were able to transposon-repeat mask 9.27% of the assembly made up of more than 500 bases (L.flavolineata.RM.min500.euk.fasta)

#### Generation of genome-guided RNASeq alignments

We aligned Illumina paired-end RNASeq data representative of *L. flavolineata* (LF_pool) tissues to the assembly of the *L. flavolineata* genome using the STAR RNASeq aligner v2.5.1b (Dobin et al. 2013) using the program's default parameters for paired-end RNASeq data.

#### Transcript assembly reconstruction of RNASeq alignments

We proceeded to assemble the RNASeq alignments generated by the STAR aligner into potential genes/transcripts by using the transcriptome reconstruction programs StringTie v1.3.3b (Pertea et al. 2015), CuffLinks v2.2.1 (Trapnell et al. 2012), CLASS v2.17 (Song & Florea 2013) and Scripture beta2 (Guttman et al. 2010) using the default parameters (for paired-end data when available) for each of these tools. The output of each of these tools was then combined and passed to the PASA (Program to Assemble Spliced Alignments, v2.0.2) pipeline (Hass et al. 2003).

#### Generation of a PASA-derived transcriptome

The PASA pipeline was used to combine the output of the transcript assembly programs into a more exhaustive transcriptome as well as to generate a TransDecoder (Haas et al. 2013)-derived ab initio program training set employed to train the ab initio programs used by the pipeline. We also provided PASA with all the 40,177 RefSeq protein-coding transcripts sequences from all organisms classified as "wasps" found in the NCBI nucleotide database (https://www.ncbi.nlm.nih.gov/nuccore). In total we provided PASA pipeline (which uses GMAP/BLAT as the alignment engines) with 224,334 transcripts which it processed into 64,583 maximal PASA transcript assemblies. PASA is set to be quite stringent;  input sequences with less than 95% identity to the genomic sequence over 95% of their length were discarded. Furthermore, PASA generates maximal assembly of spliced alignments (the transcript alignments deemed as valid are clustered based on genome mapping location and assembled into gene structures that include the maximal number of compatible transcript alignments).

#### Obtaining protein-coding ab initio/evidence-based gene predictions

##### ***Geneid (geneid+introns)* L. flavolineata*-specific gene predictions***

Geneid (Blanco et al. 2007) is an ab initio gene prediction program used to find potential protein-coding genes in anonymous genomic sequences. In the context of geneid, training basically consists of computing position weight matrices (PWMs) or Markov models of order 1 for splice sites and start codons, and deriving a model of coding DNA (generally a Markov model of order 5). Furthermore, once a preliminary species-specific matrix is obtained it is further optimized by adjusting two internal matrix parameters: the cutoff of the scores of the predicted exons (eWF) and the ratio of signal to coding statistics information to be used (oWF).

The initial training set for *L. flavolineata* was generated using one of the modules of the PASA pipeline and comprised 1,583 gene models randomly selected from within the set of 3,118 longest ORFs (more than 150 amino-acids) corresponding to complete/non-overlapping genes produced by PASA. These training-set genes were also selected because they spanned over 99% of the full-length of the annotated protein sequences of the NCBI non-redundant (NR) database (using the algorithm BLASTP, E=10-3, minimum identity=25%; NR database version of Feb. 2017). Of these gene models 80% (1,482) were used to train geneid (and most of the other ab initio tools used in this study) while the remaining 20% (371) were set aside to test the accuracy of the newly developed matrices. The 1,482 *L. flavolineata* training-set protein-coding gene models included 6,441 canonical donor splice sites/6,499 canonical acceptor sites and 1,480 start codons. These start codons were used to compute PWMs while the donor and acceptors were employed to derive Markov matrices of order 1. We also had enough coding/non-coding nucleotides to derive a Markov of order 5 as a model for the coding potential. Accuracy of the geneid parameter file was tested on an evaluation “artificial scaffold”, consisting of the 371 evaluation-set concatenated gene models with 800 nucleotides of intervening sequence between each of the genes (**Table SB1**) and subsequently used to generate genome-wide predictions.

Geneid can also use external information, such as the coordinates of known introns, to improve the accuracy of its predictions. In order to take advantage of this feature of geneid we first extracted and scored all potential introns from the "spliced junctions" (SJ) file generated by the STAR RNASeq alignment tool. This resulted in a set of 131,208 introns, of which we selected 91,599 on the basis that they overlapped with geneid predictions. To measure the accuracy of the geneid gene predictions when using intronic evidence we used STAR on the artificial scaffold to generate RNASeq alignments and then used the output SJ file to obtain all the potential intron sequences. The introns not overlapping with geneid predictions were then filtered out. Subsequently, we calculated the accuracy of geneid (+introns) on the test scaffold (excluding the mono-exonic genes), which showed an improvement in the performance of the program when using introns as evidence (**Table SB1**). The parameter file (and genome-wide intron data) were then used to obtain predictions on the entire genome assembly.

The training of geneid to obtain a parameter file for *L. flavolineata* was based on the method described to obtain a *Drosophila melanogaster* geneid parameter file (Para et al. 2000). Training was performed in a “semi-automated “ fashion by employing an in–house geneid training tool (geneidTRAINer1.1).

***GlimmerHMM* L. flavolineata*-specific gene predictions.***

We also obtained *L. flavolineata* -specific matrices for the gene prediction tool glimmerHMM (Majoros et al. 2004). GlimmerHMM is an ab initio program that is based on a Generalized Hidden Markov Model (GHMM). This program also incorporates splice site models adapted from the GeneSplicer program and a decision tree adapted from glimmmerM.

We generated the species-specific matrix for this program by using a "self-training" script obtained from the University of Maryland (http://www.cbcb.umd.edu/software/GlimmerHMM). The training was performed by following the instructions provided in http://www.cbcb.umd.edu/software/GlimmerHMM/man.shtml#training. GlimmerHMM was trained on the same set of sequences used to train the gene prediction program geneid. The new glimmerHMM matrix was evaluated on the same "artificial scaffold" used to evaluate the other ab initio gene prediction parameter files described in this text (results are shown in **Table SB1**) and subsequently used to generate glimmerHMM genome-wide predictions.

***GeneMarkES and geneMarkET* L. flavolineata*-specific gene predictions***

We generated an *L. flavolineata*-specific matrix for the gene prediction program geneMark (Lomsadze et al. 2005). The GeneMark-ES and GeneMark-ET algorithms were developed for finding protein-coding genes in eukaryotic genomes without training sets. GeneMark-ES determines species-specific gene finding parameters using a self-training algorithm based on the species of interest genomic sequence. GeneMark-ET does the same as GeneMark-ES but it can also use intron data to improve the accuracy of the predictions. We generated an *L. flavolineata*-specific matrices for this program by using the training sequence fastas generated by GeneidTrainer and following the self-training instructions found both at the program developer's website (http://opal.biology.gatech.edu/) and as indicated by Lomsadze et al. (Lomsadze et al. 2005). GeneMark-ES/ET were trained on the artificial training scaffold generated by GeneidTRAINer1.1 and geneMark-ET also used the introns (overlapping with geneid predictions) derived from the SJ file generated by STAR after aligning all-tissue RNASeq data on the artificial training scaffold. The new geneMark-ES and geneMark-ET matrices were evaluated on the same artificial scaffold used to evaluate other ab initio gene prediction parameter files used in this study (**Table SB1**). Predictions were subsequently obtained by running the programs on the entire genome of *L. flavolineata*.

***Augustus (+hints)* L. flavolineata*-specific gene predictions***

We also built an *L. flavolineata* -specific parameter file for the gene prediction program Augustus (Stanke et al. 2006). Augustus is a program that predicts genes in eukaryotic genomic sequences and that is also "re-trainable". The program is based on a Hidden Markov Model and integrates a number of known methods and sub-models. In order to obtain parameter files for Augustus we employed its own training program (http://www.molecularevolution.org/molevolfiles/exercises/augustus/training.html), and used it to estimate the optimal parameters for *L. flavolineata* given the same 1,482 species-specific genes used in training the other ab initio tools used to obtain gene predictions on these species. The resulting Augustus parameter file was evaluated on the same "artificial scaffold" consisting of the 371 concatenated gene models with 800 nucleotides of intervening sequence used to evaluate the other programs previously described (**Table SB1**), and subsequently used to generate genome-wide gene predictions.

We also took advantage of Augustus' potential to use external evidence to improve its performance. We did this by obtaining a set of predictions that used the newly developed *L. flavolineata* Augustus parameter file in combination with PASA-derived transcript evidence obtained for this species. Our strategy for taking advantage of the large set of transcripts followed the methodology described in (http://augustus.gobics.de/binaries/readme.rnaseq.html) and in an article by Stanke et al. (Stanke et al. 2006) and allowed us to obtain a higher-accuracy evidence-based set of Augustus(+hints) predictions on the *L. flavolineata* assemblies (**Table SB1**). In order for the *L. flavolineata* Augustus matrix to take advantage of the external data we first had to optimize some internal parameters of the new Augustus parameter file; the exonpart bonus for hints corresponding to PASA-evidence (“E”) was given a bonus of 1xE3. Also, for every exonpart that was not supported by the PASA evidence, the probability of the gene structure was given a “malus” or a penalization of 0.997. Furthermore, complete exons predicted by Augustus that perfectly matched the exons in the external hints were given a bonus of 1xE4. The intron bonus for (PASA) hints of source E was set to 1xE5, meaning that a predicted intron would get this bonus when being exactly as in the PASA “hint”. The Augustus parameter file plus the "hints" evidence file derived from the PASA transcriptome was also tested against the artificial scaffold as before and used to predict genes on the whole genome assembly.

##### ***SNAP*** **L. flavolineata*****-specific gene predictions***

Our final source of ab initio gene predictions to be used by the EVM combiner was obtained using the program SNAP (Korf 2004) using a *L. flavolineata*-specific matrix after training the program using a suite of self-training scripts included within the package. SNAP was developed by Ian Korf and consists of a general-purpose gene finding program that can be used both on eukaryotic and prokaryotic genomes. SNAP is an acronym for “Semi-HMM-based Nucleic Aid Parser”. SNAP was trained on the same set of sequences used to train the gene prediction program geneid. The new glimmerHMM matrix was evaluated on the same “artificial scaffold” used to evaluate the other ab initio gene prediction parameter files described in this text (results are shown in **Table SB1**) and subsequently used to generate SNAP genome-wide predictions.

##### ***Combining gene prediction data of different gene prediction programs***

Geneid (with or without introns), Augustus (with or without “hints”), glimmerHMM, geneMarkES, geneMarkET and SNAP, using their newly developed *L. flavolineata*-specific parameter files were subsequently used to predict genes on the repeat-masked assembly of this genome (L.flavolineata.RM.euk.fasta). The current *L. flavolineata* assembly is made up of 3,544 scaffolds/contigs with more than 500 bases. Given the species-specific parameter file developed for the organism in this study, geneid predicted 21,663 protein-coding genes without external evidence and 9,933 sequences when using intronic data. The ab initio tool glimmerHMM produced 31,714 gene models on the scaffolds of *L. flavolineata*. The program Augustus predicted 17,250 genes on the assembly while its evidence-based variation of Augustus(+hints) produced 19,058 predictions. The programs GeneMark-ES generated 16,115 gene models whereas GeneMark-ET (using intron evidence) produced 17,839 gene predictions. SNAP generated 24,293 predictions on the assembly of *L. flavolineata*.

The full set of TransDecoder-derived gene models generated by the training-set module of the PASA pipeline and the output of the programs above were used as input to a “combiner” (Evidence Modeler; EVM r2012-06-25; Haas et al. 2008), which was the program employed to obtain the reference annotation for this genome.

##### **EVM-based genome annotation of the *L. flavolineata* assembly by combining different sources of evidence using weights.**

A combination of the Program to Assemble Spliced Alignments (PASA v2.0.2; Haas et al. 2003) and Evidence Modeler (EVM r2012-06-25; Haas et al. 2008) were used to obtain consensus coding sequence (CDS) models using three main sources of evidence: aligned transcripts, aligned proteins, and gene predictions.

##### ***PASA transcript alignments***

The *L. flavolineata* RNA sequences processed by the PASA pipeline (v2.0.2) were obtained as briefly described below. This process resulted in PASA transcript assemblies. The transcriptome was subsequently added to the PASA database.

***Protein alignments***

In order to generate protein-alignment data for EVM all 17,084 model wasp *Nasonia vitripennis* UniProt-derived protein sequences (Feb 2017), highly curated 26,634 invertebrate SwissProt proteins (Feb 2017) and NCBI protein-coding RefSeqs (40,177 sequences; Feb 2017) classified as belonging to "wasps" were split-mapped to the *L. flavolineata* genome by using the program SPALN2 (Iwata & Gotob 2012) with Hymenoptera-specific parameters.

Furthermore, we also used the spliced-protein alignment tool exonerate (Slater & Birney 2005) to map the invertebrate SwissProt protein sequences to the scaffolds of *L. flavolineata* (**Table SB2**).

***Combining the different EVM sources***

The resulting alignments were then filtered as suggested in the EVM documentation (http://evidencemodeler.sourceforge.net/). Gene predictions were obtained as previously described and also modified as recommended (http://evidencemodeler.sourceforge.net/) and added to the EVM pipeline. We also used TransDecoder annotations generated by the PASA training-set generation module that were classified as "other predictions" in the EVM weights file.

Subsequently the transcript alignments, protein alignments and the ab initio gene models were combined into consensus CDS models by EVM using different weights. The best combination of weights (shown in **Table SB2**) were selected following the instructions contained within the EVM documentation and by previously running a "mock" EVM annotations, using a wide range of different weight files, and selecting the weights file that produced the most accurate test EVM annotation on the same “artificial scaffold” consisting of the 371 concatenated gene models with 800 nucleotides of intervening sequence previously used to evaluate all ab initio programs. Furthermore, with regard to the ab initio predictions, the weights given to each of the tools was based on the accuracy of the different programs in predicting sequences on the evaluation “artificial scaffold” for this species (**Table SB1**).

EVM built 14,365 consensus gene models (Refer to **Table SB3**). The consensus CDS models were then updated with UTRs and alternative exons through four rounds of PASA’s routine to update annotations. The resulting 19,066 transcripts were grouped into (14,095) genes, transcripts and (17,208) proteins and then a pre-selected species-specific identifier was assigned to the genes, transcripts and protein products derived from them.

***EVM consensus annotation statistics***

Finally, and as a quality control, the protein products obtained from the reference annotation of this species was aligned against either the exhaustive NCBI non-redundant (NR-201701) database using the “protein vs. protein” BLASTP “flavour” of the sequence comparison tool BLAST (E=10-2 with a minimum identity of 25%) to determine what percentage of the annotated genes matched a sequence of this large biological-sequence public databases. Results showed that 79.8% of our consensus EVM reference of this species matched an NR protein given the criteria above (**Table SB3**). Furthermore, **Table SB4** contains a wide-range of statistics obtained from the *L. flavolineata* assembly and analysis of the consensus EVM protein-coding reference annotation set obtained for this species.

| Program/**param** | **SN** | **SP** | **SNe** | **SPe** | **SNg** | **SPg** |
|---|---|---|---|---|---|---|
| EVM **lflavolineata (v7 weights)** | **0.99** | **0.94** | **0.90** | **0.85** | **0.47** | **0.49** |
| Augustus+hints **lflavolineata** | 0.96 | 0.91 | 0.82 | 0.77 | 0.30 | 0.28 |
| Geneid+intron **lflavolineata** | 0.98 | 0.94 | 0.86 | 0.81 | 0.33 | 0.33 |
| Geneid **lflavolineata** | 0.96 | 0.93 | 0.77 | 0.77 | 0.19 | 0.22 |
| Augustus **lflavolineata** | 0.93 | 0.91 | 0.75 | 0.77 | 0.16 | 0.19 |
| GeneMark-ET **lflavolineata** | 0.97 | 0.90 | 0.77 | 0.74 | 0.18 | 0.22 |
| GlimmerHMM **lflavolineata** | 0.94 | 0.91 | 0.71 | 0.71 | 0.21 | 0.17 |
| GeneMark-ES **lflavolineata** | 0.96 | 0.91 | 0.74 | 0.72 | 0.12 | 0.19 |
| SNAP **lflavolineata** | 0.91 | 0.82 | 0.68 | 0.53 | 0.09 | 0.03 |

**Table SB1.** Accuracy of gene prediction on an *L. flavolineata* “artificial scaffold” consisting of 371 concatenated *L. flavolineata* test sequences (with approximately 800 nucleotides of sequence between each of the gene models) using the *ab initio* programs geneid, augustus, glimmerHMM and geneMark-ES and SNAP with *L. flavolineata* parameter files (*i.e.* “lflavolineata”) that were built for each program given the same train-set of 1,516 gene models. The exception is geneMark-ES and geneMark-ET that were trained on the artificial training scaffold generated by GeneidTRAINer1.1. The accuracy of GeneMark-ET (using intron evidence) and that of Augustus (using RNASeq and transcript evidence i.e. “augustus+hints”) were also tested for accuracy on the same set of sequences. Geneid (geneid+introns) using introns as external evidence were also evaluated. The "optimized weights" EVM annotation of the "artificial" scaffold (**Table SB2**) was also evaluated (EVM lflavolineata). (SN & SP: sensitivity & specificity at nucleotide level; SNe & SPe: sensitivity & specificity at exon level; SNg & SPg: sensitivity & specificity at gene level).

| **Type** | **Source** | **Weight** |
|---|---|---|
| **ABINITIO_PREDICTION** | Augustus | 1 |
| **ABINITIO_PREDICTION** | AugustusHints | 1.5 |
| **ABINITIO_PREDICTION** | GlimmerHMM | 0.25 |
| **ABINITIO_PREDICTION** | GeneMark-ES | 1 |

| | | |
|---|---|---|
| **ABINITIO_PREDICTION** | geneid | 1 |
| **ABINITIO_PREDICTION** | geneid+introns | 1.5 |
| **ABINITIO_PREDICTION** | GeneMark-ET | 1 |
| **ABINITIO_PREDICTION** | SNAP | 0.5 |
| **OTHER_PREDICTION** | transdecoder | 4 |
| **PROTEIN** | SPALN2 uniprot_sprot_inv | 4 |
| **PROTEIN** | SPALN2 wasp.ncbi.prot | 3 |
| **PROTEIN** | exonerate uniprot_sprot_inv | 4 |
| **PROTEIN** | SPALN_uniprot-proteome%3AUP000002358 | 3 |
| **TRANSCRIPT** | PASA | 10 |

**Table SB2.** Weights used by EVM to create a consensus CDS model for *L. flavolineata* . (SPLAN2 against *N. vitripennis* Uniprot proteins (SPALN_uniprot-proteome%3AUP000002358); SPLAN2 against invertebrate swissprot proteins (uniprot_sprot_inv): SPALN2 against "*wasp"* NCBI Model refseq proteins (SPALN2 wasp.ncbi.prot); exonerate uniprot_sprot_inv: exonerate against invertebrate swissprot proteins.

| **Type of source of evidence** | **Number of consensus gene models supported by the type of source of evidence (% of total number of EVM reference gene models supported)** |
|---|---|
| PASA transcript alignments | 9567 (66.5947%) |
| Protein alignments | 8443 (58.7707%) |
| Protein OR PASA alignments | 10058 (70.0125%) |
| Protein OR pasa OR transdecoder | 10481 (72.957%) |
| Protein AND PASA alignments | 7952 (55.3529%) |
| only ab initio evidence (more than 2 sources) | 2982 (20.7573%) |
| only ab initio evidence (more than 3 sources) | 1855 (12.9124%) |

| | |
|---|---|
| only ab initio evidence (more than 4 sources) | 930 (6.47362%) |
| SingleEXON genes | 1960 (13.6433%) |
| singleEXON (supported by at least 2 sources of ab initio data ) | 944 (48.1633% of single exon genes) |
| singleEXON(supported by at least 2 sources of ab initio data or protein/PASA/transdecoder data ) | 1183 (60.3571% of single exon genes) |

**Table SB3.** Breakdown of the types of evidence used to build the 14,365 gene-models EVM consensus set. More than 72.9% of the gene models were built using external evidence (from protein and/or transcript data) and an additional 12.9% of the gene models were built from ab initio data that used at least three sources of evidence (*i.e.* three different programs).

| Annotation version | ***L. flavolineata 2a*** |
|---|---|
| Genome length (Mbases) | **291.28** |
| number of scaffolds | **3,544** |
| scaffolds containing annotations | **585** |
| Number of protein-coding genes | **14,095** |
| Gene density (genes/Kbase) | **0.048** |
| Number of protein-coding transcripts | **19,066** |
| Transcripts/gene (range) (% genes with more than 1 transcript) | **1.353 (SD 2.122) (1 – 163) (13.99%)** |
| Number of transcripts with UTRs | **9,588** |
| Number of proteins | **17,208** |
| Number of complete proteins (%) | **16,544 (96.14%)** |

| | |
|---|---|
| Number/(%) proteins with similarity to sequences in the NCBI NR database ($E=10^{-2}$; min. identity=25%) | **13,724 (79.75%)** |
| Avg. length of proteins | **521.691 SD 608.869** |
| Avg. length of full-length proteins | **532.795 SD 614.193** |
| Number of partial proteins (not starting with "M") | **382 (2.22%)** |
| Avg. length of partial proteins (not starting with "M") | **259.971 SD 322.42** |
| Number of partial proteins (no terminal STOP codon) | **395 (2.30%)** |
| Avg. length of partial proteins (no terminal STOP codon) | **208.678 SD 349.129** |
| Number of partial proteins (not starting with an M -and- no terminal STOP codon) | **113 (0.66%)** |
| Avg. length of partial proteins (not starting with an M -and- no terminal STOP codon) | **168.487 SD 140.218** |
| Number of partial proteins (not starting with an M -or- no terminal STOP codon) | **664 (3.86%)** |
| Avg. length of partial proteins (not starting with an M -or- no terminal STOP codon) | **245.027 SD 359.009** |
| Number of protein-coding exons | **124,103** |
| Number of introns | **105,120** |
| Number of UTRs (spliced) | **23,849** |
| Exons/transcript (range) (excludes single-exon genes) | **7.18 SD 5.86 (2 – 67)** |
| Introns/transcript (range) | **6.18 SD 5.86 (1 – 66)** |
| "spliced" UTRs/transcript (range) | **2.49 SD 0.96 (1 - 13)** |

| | |
|---|---|
| Avg. length of introns (range) | **641.706 SD 2640.1 (21 - 138002)** |
| Avg. length of mono-exonic genes | **451.57 SD 641.33** |
| Avg. length of exons (excludes mono-exonic genes) | **241.35 SD 341.65** |
| Avg. length of CDS (range) | **1,593.15 SD 1,782.17 (126 – 37,212)** |
| Avg. length of UTRs (range) | **648.92 SD 1076.59 (1 – 14,716)** |
| Avg. length of primary transcripts | **8,784.69 SD 18,960** |
| G+C content exonic (includes mono-exonic genes) | **42.3115% SD 8.8697%** |
| G+C content intronic | **28.2448% SD 10.0689%** |
| G+C content UTRs | **33.4238% SD 12.9958%** |

**Table SB4.** Statistics for the EVM-generated protein-coding annotation reference set for *L. flavolineata.*

**Associated files**

The relevant files generated by the structural annotation protocol described in this document can be found at:

**https://public-docs.crg.eu/rguigo/Data/fcamara/liastenogaster.v2a**
