## Supplemental Document S3 for "Molecular signatures of alternative fitness strategies in a facultatively social hover wasp"

**Supplementary Document S3 – Functional annotation of the *Liostenogaster flavolineata* genome**

### Methods

For the functional annotation we used InterPro (Hunter et al. 2012), KEGG (Kanehisa et al. 2012), Blast2GO (Götz et al. 2008), signalP (Petersen et al. 2011), and NCBI CDsearch (Marchler-Bauer et al. 2010) databases. InterProScan v.5.19-58 (Zdobnov & Apweiler 2001) was used to scan though all available InterPro databases, including PANTHER, Pfam, TIGRFAM, HAMAP and SUPERFAMILY. BLASTP v.2.2.29+ search against NCBI non-redundant (NR) collection of protein sequences (release 2017-06) was used as input to the local software p2gpipe version 2.5.0, database update 2017-01. KEGG orthology (KO) groups were assigned by KEGG Automatic Annotation Server (KAAS; Moriya et al. 2007) using bi-directional best hit (BBH) method against a representative gene set from 32 different species, including the mite species *Ixodes scapularis* (black-legged tick). KO identifiers were then used to retrieve using the KEGG REST-based API service the KEGG relevant functional annotation, KEGG release v.85.1.

### Results

A total of 13,137 (76.34%) out of 17,208 proteins had some type of annotation feature derived from one of the annotation resources used in this work. GO terms were assigned to 9,954(57.85%) proteins (**Table SC1**). Additionally, we were capable of assigning a description (name) to 8,167 proteins using Blast best hit or KEGG.

| | **genes** | **proteins** |
|---|---|---|
| Total number | 14,095 | 17,208 |
| **Annotated** | **10,131 (71.88%)** | **13137 (76.34%)** |
| Interpro signatures | 9,707 (68.86%) | 12,665 (73.59%) |
| Assigned to KO groups | 5,297 (37.5%) | 6,894 (40.06%) |
| With GO terms association | 7,442 (52.79%) | 9,954 (57.84%) |
| Conserved domains signatures | 8,803 (62.45%) | 11,655 (67.73%) |
| Conserved features signatures | 4,356 (30.9%) | 5,917 (34.38%) |
| SignalP signatures | 1,157 (8.20%) | 1,320 (7.67%) |

**Table SC1.** Gene and protein annotation statistics.

#### *Domain and family signatures*

In this functional annotation we used InterProScan and Batch CD-search software to assign domains and other functional elements to the proteins of interests. InterProScan v.5.19-58 was used to inspect proteins for signatures using all available InterPro databases and scanning applications, in total, 12,665 (73.59 %) proteins have some type of protein signatures. More specifically, 11,152 proteins (68.06%) are annotated with at least one signature coming from one of the most important InterPro databases for functional annotation (i.e. PANTHER, Pfam, TIGRFAM, HAMAP, SUPERFAMILY). **Table B2** displays the number of proteins containing a signature belonging to each specific InterPro member database.

Automatic Batch CD-server was used to scan a set of pre-calculated position-specific scoring matrices with proteins. In total, 11,655 proteins have domain hits and 4,356 proteins have features data. Example of annotated features: active sites, inter-domain contacts, cleavage sites or proline interaction residues.

| **InterPro member database** | **Number of proteins** |
|---|---|
| PANTHER | 11130(64.68%) |
| Pfam | 10711(62.24%) |
| SUPERFAMILY | 8775(50.99%) |
| Gene3D | 8244(47.91%) |
| ProSiteProfiles | 5841(33.94%) |
| SMART | 5257(30.55%) |
| Coils | 4370(25.40%) |
| ProSitePatterns | 3242(18.84%) |
| PRINTS | 2370(13.77%) |
| TIGRFAM | 787(4.57%) |
| PIRSF | 645(3.75%) |
| Hamap | 288(1.67%) |
| ProDom | 151(0.88%) |

**Table SC2.** Number of protein signatures identified by InterProScan for each of the InterPro member databases

***GO terms***

We have three different sources of evidence to associate GO terms to our proteins: InterPro, KEGG and p2gpipe (**Table SC3**), each of this evidence is complementary to each other. In total we managed to associate at least one GO term to 9954 proteins; with 1-24 GO terms per protein.

**Table SC4** displays the number of GO terms of each specific type obtained in this work.

| Source | Number of proteins |
|---|---|
| b2gopipe | 4033(23.44%) |
| InterPro | 9038(52.52%) |
| KEGG | 2861(16.63%) |

**Table SC3.** Source of evidence used for GO terms association and number of transcripts assigned by each of them

| Term type | Number of proteins |
|---|---|
| Molecular function | 8668 |
| Biological process | 5915 |
| Cellular component | 4308 |

**Table SC4**. Number of proteins, associated with each different GO term type.

***Proteins with identified transposon activity***

We annotated proteins as putative transposons by using annotation signatures that were previously associated to the TEs activity. Within these signatures are the PFAM domains – we're searching for ~80 transposase domains; GO terms GO:0006278(RNA-dependent DNA replication), GO:0015074(DNA integration), GO:0006355(regulation of transcription, DNA-templated), GO:0004803(transposase activity); and finally we do search for 'retrotransposon/transposase' within definition obtained from the Blast2GO or KEGG.

In total we annotate 489 proteins ( 363 genes) as putative transposons:

| Transposones annotated with | Number of proteins |
|---|---|
| PFAM domains | 37 |
| GO terms | 453 |
| Blast/KEGG definition | 3 |
| By three methods | 0 |
| By two methods | 4 |

**Table SC5.** Number of proteins associated with transposon activity.

**Associated files**

The files containing all the annotation features produced in this work can be downloaded from

https://public_docs.crg.es/rguigo/Data/avlasova/FunctionalAnnotation/waspProject/L.flavolineata
