## Supplemental Figures S1-2 for "Molecular signatures of alternative fitness strategies in a facultatively social hover wasp"

**Figure S1.** WGCNA summary network measures against soft thresholding power. Numbers in the plots indicate the corresponding soft thresholding powers.

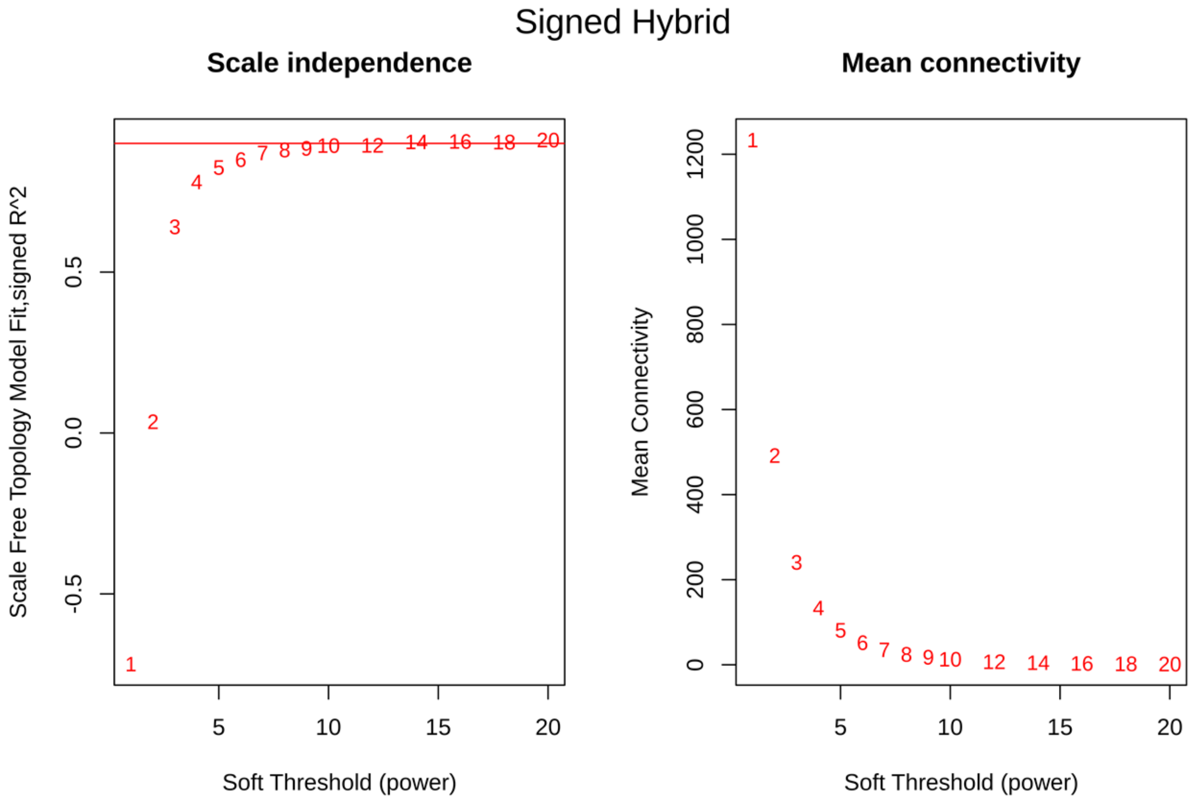

**Figure S2.** Gene dendrogram with clustering based on consensus topological overlap. Upper colour row: consensus module assignments prior to merging of modules with similar expression profiles. Lower colour row: consensus modules following merging of similar modules.

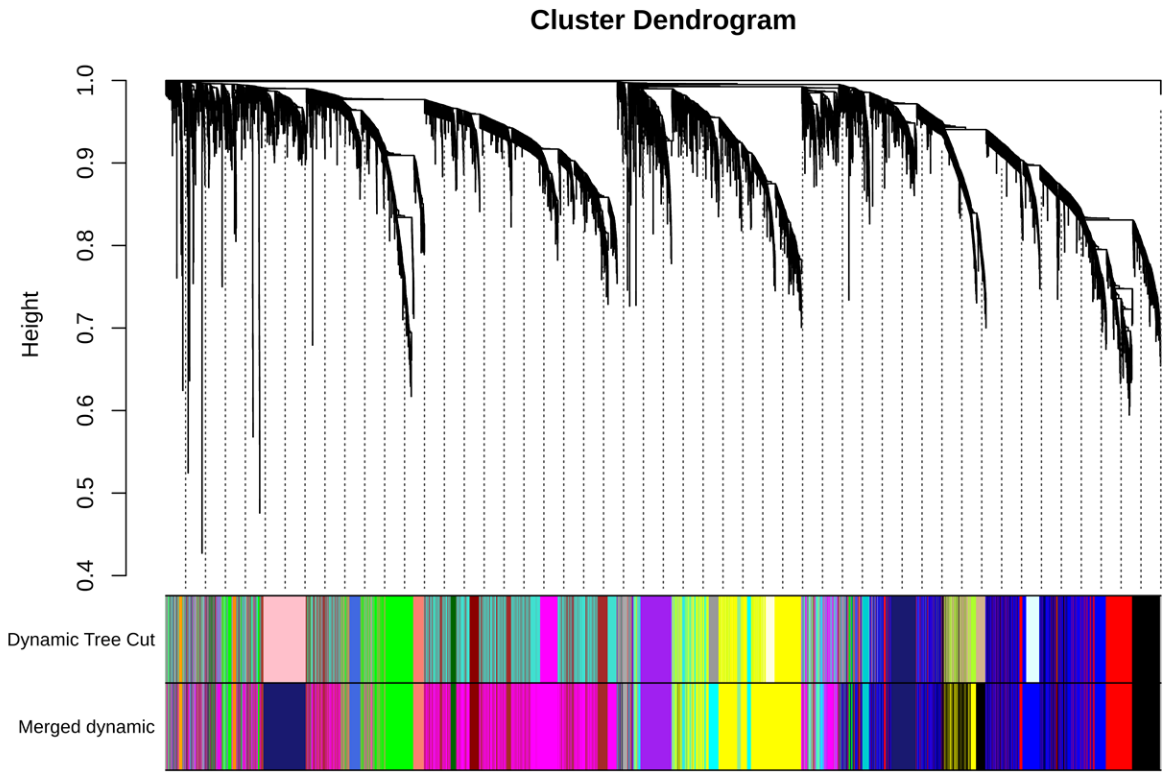
